# IMP: a pipeline for reproducible integrated metagenomic and metatranscriptomic analyses

**DOI:** 10.1101/039263

**Authors:** Shaman Narayanasamy, Yohan Jarosz, Emilie E.L. Muller, Cédric C. Laczny, Malte Herold, Anne Kaysen, Anna Heintz-Buschart, Nicolás Pinel, Patrick May, Paul Wilmes

## Abstract

We present IMP, an automated pipeline for reproducible integrated analyses of coupled metagenomic and metatranscriptomic data. IMP incorporates preprocessing, iterative co-assembly of metagenomic and metatranscriptomic data, analyses of microbial community structure and function as well as genomic signature-based visualizations. Complementary use of metagenomic and metatranscriptomic data improves assembly quality and enables the estimation of both population abundance and community activity while allowing the recovery and analysis of potentially important components, such as RNA viruses. IMP is containerized using Docker which ensures reproducibility. IMP is available at http://r3lab.uni.lu/web/imp/.

## Background and motivation

Microbial communities are ubiquitous in nature and govern important processes related to human health and biotechnology [1, 2]. A significant fraction of naturally occurring microorganisms elude detection and investigation using classical microbiological methods due to their unculturability under standard laboratory conditions [3]. The issue of unculturability is largely circumvented through the direct application of high-resolution and high-throughput molecular measurements, most notably metagenomics, to microbial community samples collected *in situ* [4–6]. Beyond metagenomics, there is also a clear need to obtain functional readouts in the form of additional layers of omics data from consortia. Moreover, there is a growing desire to integrate the resulting meta-omics data to more conclusively link genetic potential to actual phenotype *in situ* [6]. For this purpose, specialized wet-lab methods have been developed to ensure that the generated data fulfill the premise of systematic measurements [7], as subsampling has been shown to inflate intra- and inter-sample variation, thereby hampering subsequent data integration, individual biomolecular fractions, i.e. DNA, RNA, proteins and metabolites are derived from single, unique samples [7, 8]. Next-generation sequencing (NGS) of microbial community derived DNA and reverse transcribed RNA (cDNA) results in metagenomic (MG) and metatranscriptomic (MT) data, respectively. Additional layers of meta-omic data include the metaproteome (MP) and the (meta-)metabolome [4–6, 9]. Although standardized and reproducible wet-lab methods have been developed for integrated omics of microbial communities, corresponding dry-lab workflows have yet to be formalized.

Computational solutions for the analysis of MG and MT data can be broadly categorized into reference-dependent or reference-independent (*de novo*) methods [5]. Reference-dependent methods are based on the direct alignment/mapping of sequencing reads onto isolate genomes, gene catalogs or MG data. A major drawback of such methods is the large number of sequencing reads discarded during data analysis, due to their dissimilarity from genomes/genes within the reference databases. More specifically, based on analyses of MG data from the human gut microbiome, which is arguably the most resolved microbial ecosystem in terms of representative isolate genomes, 43% of organisms are not represented by isolate genomes [10], while 74%-81% of sequencing data is typically represented within an integrated gene catalog based on MG data [11], exemplifying a substantial loss of potentially useful information when using isolate genome reference databases alone. Conversely, reference-independent methodologies, such as approaches based on *de novo* assembly, enable the retrieval of previously uncharacterized genomes and/or potentially novel genes, providing an added advantage over reference-based methods [4, 5, 12]. Furthermore, it has been shown that the assembly of sequencing reads into longer contiguous sequences (contigs) greatly improves the taxonomic assignments and prediction of genes as opposed to direct identification from short sequencing reads [13, 14].

Given the advantages of reference-independent methods, a wide array of MG-specific assemblers such as IDBA-UD [15] and MEGAHIT [16] have been developed. Most MT analyses consist of reference genome- or metagenome-dependent workflows, similar to the reference-dependent workflows used for MG data [17, 18]. However, reference-independent approaches for metatranscriptomic data exploitation are also available either using specialized metatranscriptome assemblers (e.g. IDBA-MT [19]), metagenome assemblers (e.g. IDBA-UD [15]) or single-species transcriptome assemblers (e.g. Trinity [20]) [14]. In both cases, the available assemblers are able to handle the uneven sequencing depths of MG and MT data. Although dedicated assembly methods have been developed for MG and MT data, formalized pipelines allowing both data types to be used in an integrated way have yet to be developed.

Automated data processing and analysis pipelines have so far been mainly developed for MG data. These include pipelines such as MOCAT [21] and MetAMOS [12] which incorporate the entire process of MG data analysis, ranging from preprocessing of sequencing reads to *de novo* assembly and post-assembly analysis (read alignment, taxonomic classification, gene annotation, etc.). Both pipelines use SOAPdenovo [22] as the default *de novo* assembler, performing single-length *k*mer-based assemblies which usually result in fragmented (low contiguity) assemblies with lower coverage levels, compared to multiple-length *k*mer-based assemblers [23]. However, MetAMOS offers more flexibility by providing multiple options for MG assemblers, in addition to being an easily installable and user-amendable pipeline for standardized MG data analysis. *De novo* MG assemblies may be further leveraged by binning the data to resolve and retrieve population-level genomes including those from hitherto undescribed taxa [17, 24–28].

Multi-omic analyses have already allowed unprecedented insights into microbial community structure and function *in situ,* using different *in silico* analysis approaches. Franzosa *et al.* (2014) [29] have applied reference-based analyses of MG and MT data to study healthy human fecal microbial community samples. Conversely, Hultman *et al.* (2015) [30] performed a multi-omic survey of microbial communities in permafrost soils, leveraging coupled MG, MT and MP data. The MG data was subjected to *de novo* assembly, annotation and binning of the resulting contigs. The remaining MT and MP datasets were analyzed based on the MG assembly and additional reference databases. Bremges *et al.* (2015) [31] performed analysis on a production-scale bioreactor, using deep MG and MT data. *De novo* assembly and annotation was carried out for the MG data, after which MT data was used to identify active metabolic pathways of target organisms. Furthermore, the authors bundled all tools and dependencies within a Docker container to promote reproducibility of their workflow. In contrast to these aforementioned studies, arguably the most integrated multi-omic study to date was performed by Muller, Pinel *et al.* (2014) [32], which involved a temporal survey of a oleaginous microbial community from a biological wastewater treatment plant using a combination of MG, MT and MP data. In their study, MG and MT data were integrated via a *de novo* co-assembly procedure, the output of which was subsequently binned and annotated. The resulting gene set was subsequently used to identify peptides from the MP data. In addition, patterns of intra- and inter-population expression and genomic variation (single nucleotide polymorphisms, SNPs) were resolved. Furthermore, Roume, Heintz-Buschart *et al.* (2015) [33] performed a comparative study of two samples from a biological wastewater treatment plant. In this work, MG and MT data were also integrated through co-assemblies and the MP data was analyzed in relation to these co-assemblies. The latter two studies performed integration by co-assembling MG and MT data. Although these studies clearly demonstrated the power of multi-omic analyses in facilitating unprecedented insights into community structure and function, standardized and reproducible dry-lab workflows for integrating and analyzing the multi-omic data have so far been unavailable. Such approaches are pertinent to compare results between different studies and systems of study.

While MG analysis provides information on the gene coding potential (functional potential) of a given community, complementary usage of MT data enables the study of transcriptional activity, which more faithfully represents potential community-wide phenotypes [4, 9, 34]. Due to the absence of tools/workflows to handle multi-omic datasets, most of the aforementioned studies utilize non-standardized, *ad hoc* analyses, mostly made up of custom workflows, creating a challenge in reproducing the analyses [12, 35–37]. Here, we present the Integrated Meta-omic Pipeline (IMP), an open source *de novo* assembly-based pipeline to perform standardized, automated and reproducible large-scale integrated analysis of multi-omic (MG and MT) data, derived from a single microbial community.

## Overview of IMP

IMP leverages Docker for reproducibility and deployment. The interfacing with Docker is facilitated through a user-friendly Python wrapper script. As such, Python and Docker are the only prerequisites for the pipeline, enabling an easy installation and execution process. Workflow implementation and automation is achieved using Snakemake [38, 39]. The IMP workflow can be broadly categorized into three major processes: i) preprocessing, ii) assembly and iii) analysis (Fig. 1). The modular design and open source features of IMP also allow for customization of the pipeline to suit specific user-defined analysis requirements. Detailed parameters for IMP processes are described in the section Details and parameters of IMP and examples of detailed workflow schematic is provided in Additional file 1: HTML S1 and S2.

**Fig. 1.**
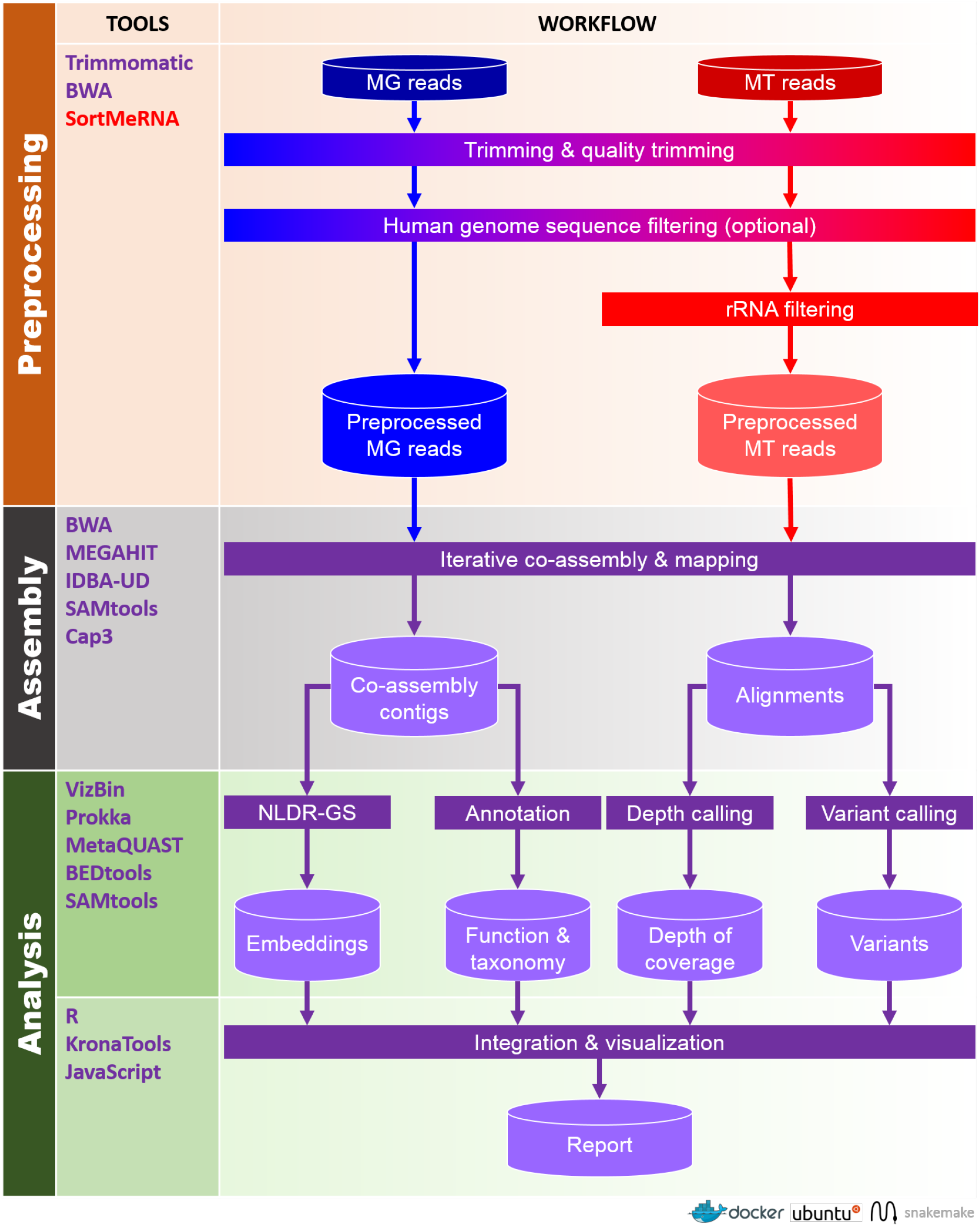
Schematic overview of the IMP pipeline. Cylinders represent input and output while rectangles represent processes. MG: Metagenomic data, MT: Metatranscriptomic data, rRNA: ribosomal RNA and NLDR-GS: non-linear dimensionality reduction of genomic signatures. IMP handles raw paired-end (unprocessed) MG and MT data. Processes, input and output specific to MG and MT data are labeled blue and red, respectively. Processes and output that involve integration of MG and MT data are represented in purple. Details about the “iterative co-assembly” are available in Additional file 2: Figure S1. IMP is launched as Docker container with Ubuntu as the operating system and uses Snakemake for workflow automation.

### Preprocessing of paired-end reads

The input to IMP consists of MG and MT (preferably depleted of ribosomal RNA - rRNA) paired-end reads in FASTQ format, each comprising a set of two files containing pair-1 and pair-2 reads, respectively. MG and MT reads are preprocessed independently of each other and this involves an initial quality control step (see Fig. 1 and section Trimming and quality filtering) [40] followed by filtering of reads that are deemed undesired in downstream processes. For the analysis of MG and MT data from human microbiome studies, the quality control steps should be followed by the optional filtering of human genome derived sequences (see Fig. 1 and section Human genome derived sequence filtering). Additionally, *in silico* rRNA sequence depletion is applied to the MT data (see Fig. 1 and section Ribosomal RNA filtering). In summary, the preprocessing implemented in IMP involves the independent removal of sequences from MG and MT datasets which are undesired due to technical and/or biological reasons.

### IMP-based iterative co-assembly

IMP implements an iterative co-assembly procedure (Fig. 1 and Additional file 2: Figure S1), which combines the benefits from iterative assemblies and co-assemblies of multi-omic (MG and MT) data [32]. Additional rounds of assembly involving the unused reads (unmappable) from previous assembly are herein referred to as “iterative assemblies”, while assemblies involving both MG and MT reads are hereafter referred to as “co-assemblies”.

Preprocessed MT reads are first assembled to generate an initial set of MT contigs (Additional file 2: Figure S1). MT reads that remain unmapped to the initial set of MT contigs undergo an additional round of assembly. The combined set of MT contigs (from both the aforementioned MT assemblies) are then used as input, together with preprocessed MG and MT reads, to perform an initial co-assembly. The MG and MT reads which remain unmapped to the resulting contigs are recruited for an additional co-assembly iteration. The resulting contigs are further refined by performing a contig-level assembly, which aligns highly similar contigs against each other. This procedure aims at reducing redundancy by collapsing shorter contigs into longer contigs and/or improving contiguity by extending contigs via overlapping contig ends, thereby producing the final set of contigs (Additional file 2: Figure S1). The IMP-based iterative co-assembly is a key feature that facilitates the integration of MG and MT data as it allows maximization of overall data usage. Finally, preprocessed MG and MT reads are mapped onto the final contigs and the resulting alignment information is used in various downstream analysis procedures (Fig. 1). Please refer to section Details and parameters of IMP for information about programs and parameters.

### Post-assembly analysis and output

The set of contigs resulting from the IMP-based iterative co-assembly undergo quality assessment as well as taxonomic [41] and functional annotation [42]. Additionally, non-linear dimensionality reduction of genomic signatures (NLDR-GS) is performed using VizBin [24, 43] which provides two-dimensional (2D) embeddings, enabling the visualization of the contigs as scatter plots in a 2D map format. Further analysis steps include, but are not limited to, calculations of the contig- and gene-level depths of coverage and the calling of genomic variants (using three variant callers, see section Variant calling). The information from these analyses are condensed and integrated into the VizBin-based maps to produce augmented visualizations. The visualizations and various summaries of the output are compiled into a HTML report (for examples of HTML report see Additional file 1: HTML S1 and S2 and for details of output see section Output).

## Results and discussion

We demonstrate the performance and output of IMP on three multi-omic datasets, each consisting of MG and MT paired-end reads (see section Coupled metagenomic and metatranscriptomic datasets for details). A simulated mock (SM) community of 73 bacterial genomes [14] was mainly used to benchmark the IMP-based iterative co-assemblies in comparison to standard assembly strategies. Additional benchmarking of the iterative co-assemblies were performed on published datasets from a human fecal (HF) sample [29] and a wastewater (WW) sludge microbial community [32]. The latter datasets were used to assess the output and features of IMP ranging from preprocessing to post-assembly analyses.

### Preprocessing and filtering

The preprocessing and filtering of sequencing reads is essential for the removal of low quality bases/reads and potentially unwanted sequences, prior to assembly and analysis. Preprocessing of NGS data, prior to assembly, has been shown to increase the quality of *de novo* assemblies, despite decreased numbers of input reads [44]. The preprocessing of MG and MT reads is handled in a tailored manner within IMP.

The results of the IMP and MetAMOS preprocessing procedures are summarized in Additional file 3: Table S1. The preprocessing of the HF included the optional filtering of human genome derived-sequences, while the same step was omitted for the WW data. Even though the preprocessing using IMP yields both paired- and single-end reads, the following section discusses only the paired-end reads as they make up a large fraction of the data and are generally more informative compared to single-end reads. However, IMP retains the use of single-end reads in all downstream processes, unlike most available methods, which discard such reads. Since MetAMOS assumes all input to be MG data, it cannot be directly compared against the preprocessing using IMP.

The final output of IMP’s preprocessing and filtering procedure retained 69.2% (29.8% low quality and 1.0% human genome derived sequences) and 89.2% (10.8% low quality) of MG paired-end reads for HF and WW, respectively. Similarly, approximately 90.8% (6.4% low quality, 1.2% rRNA and 1.7% human genome derived sequences) and 55.3% (19.8% low quality and 24.9% rRNA) of MT paired-end reads were retained from HF and WW, respectively. The filtering of human genome derived sequences are due to technical and privacy reasons, whereas *in silico* rRNA filtering helps remove remaining rRNA reads, which are usually abundant in cDNA libraries, even after the application of wet-lab rRNA depletion procedures. In summary, IMP is designed to perform stringent and standardized preprocessing of MG and MT data in a tailored way enabling efficient data usage in subsequent steps.

### Assessment of the iterative assembly approach

*De novo* assemblies of MG or MT data usually result in a large fraction of reads that are unmappable to the produced contigs and therefore remain unused, resulting in suboptimal data usage. Previous studies (e.g. Muller, Pinel *et al.* 2014 [32]; Schürch *et al.* 2014 [45]; Reyes *et al.* 2015 [46]) assembled the set of unmappable reads iteratively to successfully obtain additional contigs from these additional rounds of assembly and which lead to an increase in number of predictable genes.

In order to evaluate the best iterative assembly approach for IMP, we attempted to determine the opportune number of assembly iterations in relation to assembly quality metrics. The evaluation involved performing multiple iterations of recruiting unmappable reads to the previously generated non-redundant assembly, followed by a *de novo* assembly of those unmapped reads. The assembly from a given iteration was then merged with the previous assembly to reduce the redundancy (refer to section Iterative single-omic assemblies for details). The evaluation of additional assembly iterations for MG data of SM, HF and WW are summarized in Fig. 2, based on four different metrics. Overall, each iteration on each of the different datasets (SM, HF and WW) lead to an increase in total length of assembly and increased the overall number of mappable reads, but differed in the observed gain of contigs and genes (Fig. 2; and Additional file 3: Table S2). A similar trend is noticeable for iterative assemblies on MT data (see Additional file 2: Figure S2 and Additional file 3: Table S3). The observed trends may be explained by the fact that the complexity of the data typically confounds assemblies [44]. The exclusion of mappable reads in each iteration of assembly reduces the complexity of the data, which in turn allows additional contigs to be assembled and results in a higher cumulative output [44].

**Fig. 2.**
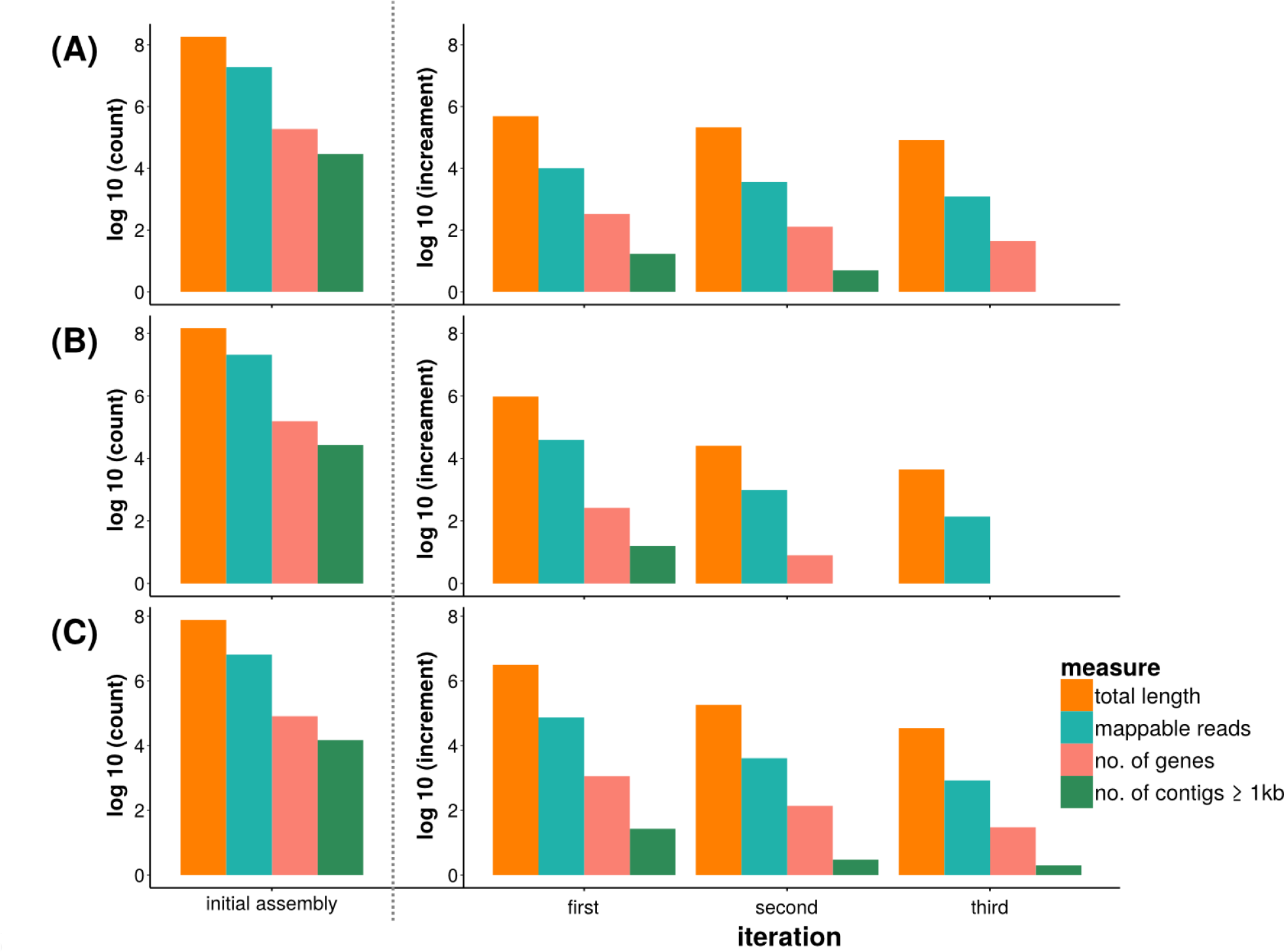
Information contained within the iterative metagenomic assemblies. Quantitative assessment of the initial metagenomic (MG) assembly as well as incremental information and data usage from additional MG assembly iterations, employing unmappable reads from: **(A)** simulated mock (SM) community, **(B)** human fecal (HF) community and **(C)** wastewater (WW) sludge community.

Considering the relatively low increase in longer contigs and genes beyond the first assembly iteration (Fig. 2, Additional file 2: Figure S2 and Additional file 3: Table S2 and S3) and the extended runtimes required to perform additional assembly iterations, an opportune single iteration assembly approach was implemented in the workflow of IMP. This approach, which balances the maximization of output yield with runtime, is implemented within the IMP-based co-assembly approach.

### Benchmarking

The iterative co-assemblies were benchmarked against single-omic MG and MT assemblies and co-assemblies obtained using the state-of-the-art MG data analysis pipeline, MetAMOS [12].

#### Single-omic assemblies and multi-omic iterative co-assemblies

Separate single-omic iterative assemblies on all datasets were generated using the preprocessed MG and MT data (see Iterative single-omic assemblies). The iterative co-assemblies were executed in IMP using the two available assembler options for the co-assembly step, i.e. the default IDBA-UD [15] (hereafter referred to as IMP) and the optional MEGAHIT assembler [16] (referred to as IMP-MEGAHIT). Both assemblers are regarded as state-of-the-art because they perform assemblies on multiple *k*mer sizes, while MEGAHIT was also chosen due to its superior speed and efficient memory usage [15, 16].

The comparison between single-omic assemblies and IMP-based co-assemblies are summarized in Table 1. The IMP-based co-assemblies consistently returned larger number of contigs, increased total length of assembled contigs and a higher number of predicted genes (partial genes included) compared to single-omic assemblies for all datasets (Table 1). The apparent slight reduction in contiguity (N50 statistic) is due to the addition of shorter contigs likely stemming from the increased sequencing depth of the combined MG and MT datasets, which also increases the complexity of the assembly process. By using the reference genomes from the SM data as ground truth, an improved recovery of reference genome fractions is apparent for the IMP-based co-assemblies. Importantly, a significant increase in the number of mappable MG and MT reads was observed within all co-assemblies compared to the respective single-omic assemblies (Table 1) which suggests superior data usage using the IMP-based approach. For example, the IMP-based iterative co-assemblies resulted in a large fraction of reads being mappable back to the contigs derived from the HF sample (average of approximately 88.0 % and 96.3 % for the MG and MT reads, respectively; Table 1), which is substantially higher compared to the numbers reported in a previous report in which MG sequencing data was mapped to an integrated gene catalog, i.e. 74%-81% [11]. In summary, the complementary use of MG and MT data in the context of *de novo* co-assembly results in an increased yield of output, while enhancing overall data usage for subsequent analyses.

**Table 1.** Co-assemblies versus separate single-omic assemblies. Characteristics of the metagenomic (MG) and metatranscriptomic (MT) co-assemblies (IMP and IMP-MEGAHIT) against MG-only and MT-only assemblies based on simulated mock (SM) community dataset, human fecal (HF) community dataset and wastewater (WW) sludge community dataset. N50 statistics are reported based on a 500bp cut-off while read mappings were performed on all contigs.

| Sample | Assembly | No. of contigs (all) | Total length | N50 | No. of predicted genes (unique) | Genome fraction (%) | MG mapped reads (%) | MT mapped reads (%) |
| --- | --- | --- | --- | --- | --- | --- | --- | --- |
| SM | IMP | 84052 | 198407791 | 10154 | 201480 | 65.3 | 97.6 | 90.2 |
|  | IMP-MEGAHIT | 108011 | 208588327 | 8243 | 212669 | 70 | 98 | 91.39 |
|  | MG-only | 74498 | 183252937 | 10902 | 187382 | 60.8 | 96.39 | 80.36 |
|  | MT-only | 14723 | 9497250 | 961 | 7357 | 3.5 | 8.58 | 29.03 |
| HF | IMP | 131728 | 174193511 | 4227 | 182924 | NA | 87.5 | 96.8 |
|  | IMP-MEGAHIT | 149163 | 181987482 | 3696 | 191503 | NA | 88 | 95.7 |
|  | MG-only | 121018 | 146897961 | 3564 | 155068 | NA | 35.1 | 34.35 |
|  | MT-only | 43466 | 46274315 | 2248 | 48527 | NA | 4.27 | 4.73 |
| WW | IMP | 167295 | 125102684 | 1501 | 126748 | NA | 30.9 | 61.6 |
|  | IMP-MEGAHIT | 208345 | 143114883 | 1285 | 143354 | NA | 32.1 | 62.8 |
|  | MG-only | 88818 | 77058077 | 2100 | 81084 | NA | 0.73 | 12.2 |
|  | MT-only | 47237 | 23251573 | 808 | 17851 | NA | 0.05 | 0.53 |

#### Quality assessment of the IMP-based iterative co-assembly procedure

Iterative co-assemblies using IMP (referred to as IMP and IMP-MEGAHIT, see section above) and co-assemblies from MetAMOS [12] on all datasets were compared against each other. MetAMOS was chosen due to its similar aim of providing an open source, reproducible and standardized *de novo* assembly-based platform for large-scale microbiome sequencing analyses [12]. Although MetAMOS was developed specifically for MG data analysis, it was hereby extended to perform MG and MT co-assemblies by including both MG and MT read libraries as input (see section Execution of pipelines) using two available assembler options: SOAPdenovo [22] (hereafter referred to as MetAMOS) and IDBA-UD [15] (hereafter referred as MetAMOS-IDBA-UD). The results of the comparison are summarized in Fig. 3 (see Additional file 2: Figure S3 and Additional file 3: Table S4 for detailed comparison and results).

**Fig. 3.**
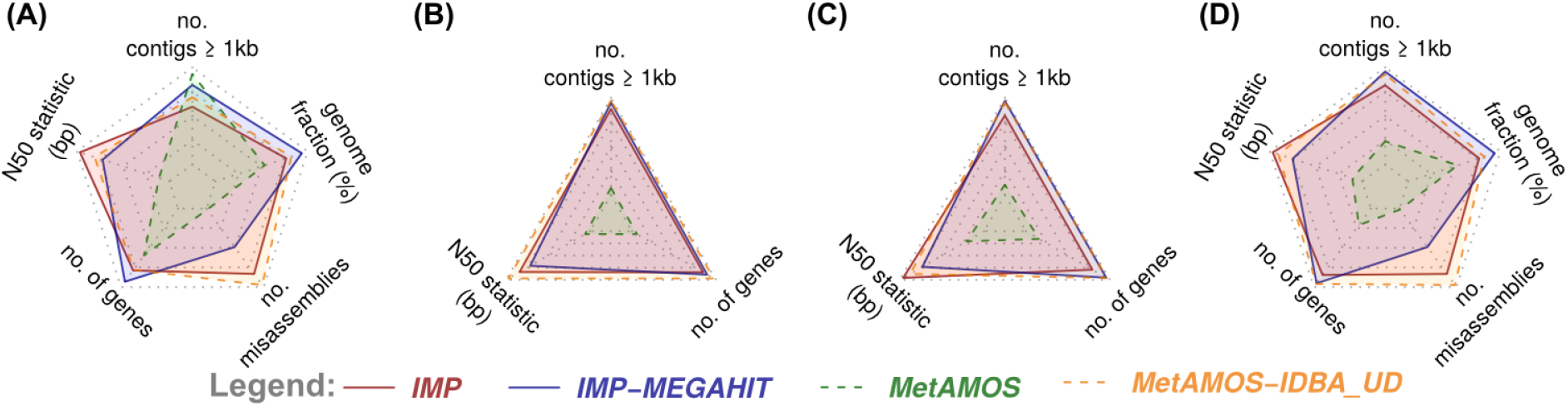
Assessment of the IMP-based iterative co-assemblies in comparison to established methods. Radar charts summarizing the characteristics of the co-assemblies generated using IMP and MetAMOS pipelines on: **(A)** simulated mock (SM) community **(B)** human fecal (HF) community and **(C)** wastewater (WW) sludge community. **(D)** Summary radar chart reflecting the cumulative measures obtained using the different datasets. The solid lines represent IMP assemblies while the dashed lines represent MetAMOS assemblies, both pipelines executed using two different assemblers. The assemblies are assessed based on number of contigs ≥ 1kb, N50 statistics (contiguity), number of predicted genes (unique). N50 statistics are reported using a 500bp cut-off. Additional reference-based assessments for SM assemblies include recovered genome fraction (%) and proportion of misassemblies. Higher values within the radar charts (furthest from center) represent best performance, except for misassemblies, where lower (closer to center) values indicate best performance.

Based on the SM data (Fig. 3A), IMP and MetAMOS-IDBA-UD performed similarly for most measures but the IMP-based assemblies producing slightly better contiguity (N50 statistic) and lower levels of apparent misassembly. Despite MetAMOS producing the largest number of contigs ≥ 1kb, its comparatively low N50 statistic indicates a highly fragmented assembly, which is further reflected in the low number of predicted unique genes. Conversely, this fragmented assembly is accompanied by relatively low misassembly rate, reinforcing the notion that shorter contigs are less prone to misassemblies [44]. However, longer contigs (≥ 1kb) are a prerequisite for population-level genome reconstruction and subsequent multi-omic data interpretation. On the other hand, IMP-MEGAHIT generated the highest number of predicted unique genes, recovered the largest fraction of reference genomes, while yielding comparatively large number of contigs ≥ 1kb, relatively high N50 statistic and a low rate of misassembly. The assessment based on real datasets shows comparable performance between IMP, IMP-MEGAHIT and MetAMOS-IDBA_UD (Fig. 3B and C). In general, MetAMOS produced highly fragmented assemblies for the real datasets, possibly due to the single *k*mer length (*k* = 31) assemblies, which tend to produce relatively fragmented assemblies compared to multiple *k*mer length assemblers (as in IMP, IMP-MEGAHIT and MetAMOS-IDBA-UD) [23]. In summary (Fig. 3D), IMP and MetAMOS-IDBA-UD performed similarly for most metrics when the same assembly program (IDBA-UD) was used by both pipelines. However, the IMP iterative co-assemblies were generated using a lower number of reads compared to MetAMOS-IDBA_UD due to the more stringent preprocessing procedures in IMP, which in turn yielded better quality assemblies (Fig. 3D) which are a prerequisite for population-level genome reconstruction and multi-omic data interpretation.

#### Summary output from IMP

The workflow of IMP is unique such that it allows integrated MG and MT data handling. Although MetAMOS may be extended to perform co-assemblies of MG and MT data, it does not discriminate between the two data types in its pre- and post-assembly procedures which is important given the disparate nature of MG and MT datasets.

IMP generates several output files, as detailed in the Output section which allow both reference-dependent and-independent analyses of the data. Information from these output files are condensed and summarized using different static and dynamic visualization methods, which are compiled into an HTML report (Additional file 1 : HTML S1 and S2). Fig. 4 presents selected output available from the analysis of IMP on the HF data. The generated taxonomic overviews are based on the alignment of contigs to the most closely related prokaryotic genomes within the NCBI genome database and the fraction of potential reference genome bases covered (Fig. 4A; Additional file 1: HTML S1) [41]. The abundances of the predicted genes (based on average depths of coverage) may be represented both at the MG and MT levels and thus enable the comparison of functional potential (Fig. 4B) and actual expression (Fig. 4C) of various KEGG Ontology categories (for details, see Krona charts within Additional file 1: HTML S1). IMP also provides augmented VizBin-based 2D maps [24, 43], with additional layers of information integrated onto them, for example, variant densities (Fig. 4D) and MT to MG depth of coverage ratios (Fig. 4E). These visualizations may aid users in highlighting subsets of contigs based on certain characteristics of interest, i.e. population heterogeneity/homogeneity, low/high transcriptional activity, etc. Please refer to Additional file 1: HTML S1 and S2 for further examples.

**Fig. 4.**
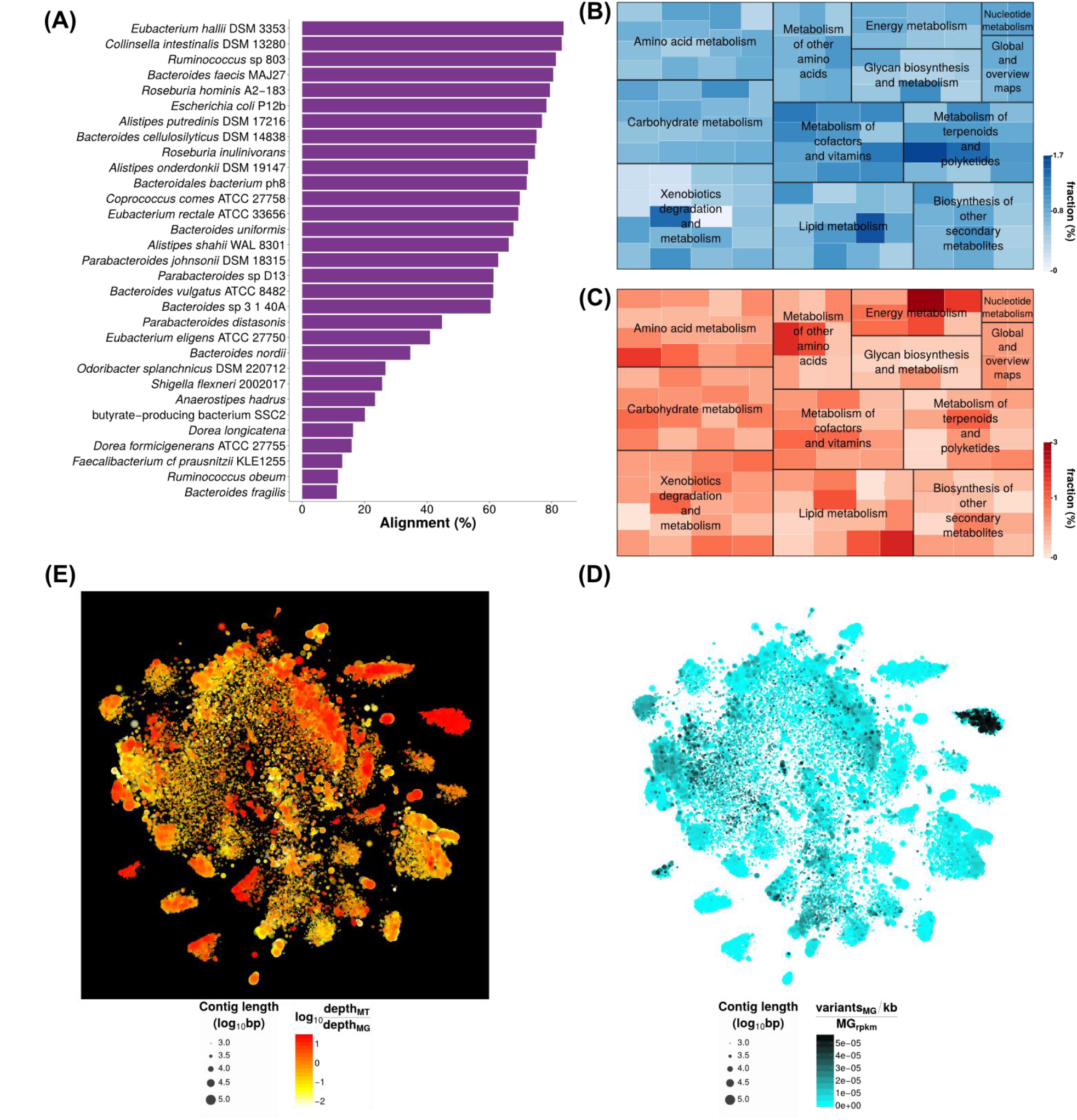
Examples of information retrievable from the IMP output of human fecal metagenomic and metatranscriptomic data. **(A)** Taxonomic composition reflecting the percentages of genomes covered. Representation of the inferred abundances of certain metabolic pathways based on **(B)** metagenomic data (functional potential) and **(C)** metatranscriptomic data (gene expression). Augmented VizBin maps of contigs ≥ 1kb, representing **(D)** contig-level metagenomic variant densities and (E) contig-level ratios of MT to MG average depth of coverage.

### Integrated omic data allows identification of key microbiome characteristics

The integration of MG and MT data provides unique opportunities for uncovering community-or population-specific traits which cannot be resolved from MG data alone. Here we provide two examples of insights gained through the direct comparison of the MG and MT results provided by IMP.

#### Identification of RNA viruses

To identify differences in the information content of MG and MT complements, the contigs generated from IMP were inspected with respect to coverage by MG and MT reads. In the two exemplary datasets, a large fraction of the contigs resulted from the composite assembly of MG and MT data, followed by contigs composed exclusively of MG data and a small proportion composed exclusively of MT data (Additional file 3: Table S5). Longer contigs (≥ 1 kb) composed exclusively of MT reads and annotated with known viral/bacteriophage genes were enriched and retained for further inspection Table 2 (for the complete list contigs, see Additional file 3: Table S6 and S7). A sequence alignment-based search against the NCBInr nucleotide database revealed that the longer contigs represent almost complete genomes of RNA viruses (see Additional file 3 : Table S8 and S9). This demonstrates that the incorporation of MT data in the assembly enables the recovery of nearly complete RNA virus genomes, thereby allowing their detailed study.

**Table 2.** Contigs reconstructed from the metatranscriptomic data with a likely viral/bacteriophage origin/function. Contigs of ≥ 1kb and average depth of coverage ≥ 20 were selected.

| Sample | Contig ID | Contig length | Avg. contig depth of coverage | Gene product | Avg. gene depth of coverage |
| --- | --- | --- | --- | --- | --- |
| HF | contig_34 | 6468 | 20926.50 | Virus coat protein (TMV like) | 30668.30 |
|  |  |  |  | Viral movement protein (MP) | 26043.40 |
|  |  |  |  | RNA dependent RNA polymerase | 22578.40 |
|  |  |  |  | Viral methyltransferase | 18816.60 |
|  | contig_13948 | 2074 | 46.00 | RNA dependent RNA polymerase | 40.98 |
|  |  |  |  | Viral movement protein (MP) | 55.76 |
| WW | contig_6405 | 4062 | 46.24 | Tombusvirus p33 | 43.42 |
|  |  |  |  | Viral RNA dependent RNA polymerase | 41.76 |
|  |  |  |  | Viral coat protein (S domain) | 36.48 |
|  | contig_7409 | 3217 | 20.62 | Viral RNA dependent RNA polymerase | 18.24 |
|  |  |  |  | Viral coat protein (S domain) | 20.88 |
|  | contig_7872 | 2955 | 77.01 | hypothetical protein | 112.11 |
|  |  |  |  | Phage maturation protein | 103.02 |

#### Identification of populations with apparent high transcriptional activity

To further demonstrate the unique analytical capabilities of IMP, we set out to identify populations with a high transcriptional activity in the HF sample. Average depth of coverage at the contig- and gene-level is a common measure used to evaluate the abundance of microbial populations within communities [25, 27, 32]. Integrative analysis of MG and MT data by IMP further extends this measure by calculation of average MT to MG depth of coverage ratios, which provides information on transcriptional activity and can be visualized using augmented VizBin maps.

One particular cluster of contigs within the augmented VizBin maps displayed high MT to MG depth of coverage ratios (see Additional file 2: Figure S4). The subset of contigs (simplified as subset) within the selected cluster aligned to the genome of the *Escherichia coli* P12B. For comparison, we also identified a subset which was highly abundant at the MG level (lower MT to MG ratio), which aligned to the genome of *Collinsella intestinalis* DSM 13280 strain. Based on these observations, we highlighted these subsets of contigs to produce an augmented VizBin map (Fig. 5A). In these, *C. intestinalis* and *E. coli* subsets are mainly represented by clear peripheral clusters which exhibit consistent intra-cluster MT to MG depth of coverage ratios (Fig. 5A). The subsets were manually inspected in terms of their distribution of average MG and MT depths of coverage, comparing them against the corresponding distributions of all the contigs. The MG-based average depths of coverage of the contigs from the entire community exhibited a bell-shape like distribution, with a clear peak (mode). On the contrary, MT depths of coverage exhibited a spread distribution, with a relatively lower mean (compared to MG distribution) and no clear peak (Fig. 5B). The *C. intestinalis* subset displays similar distributions to that of the entire community, whereas the *E. coli* subset exhibits an unusually high MT-based depth of coverage, and a low MG-based depth of coverage (Fig. 5B). Further inspection of the individual omic datasets revealed that the *E. coli* subset was not covered by the MG-based contigs, while 70 % of the *E. coli* genome was recoverable from the MT-based assembly (Fig.5C). In contrast, the *C. intestinalis* subset demonstrated comparable genomic recovery in all co-assemblies and the MG-only assembly. As noted by the authors of the original study by Franzosa *et al.* (2014) [29], the cDNA conversion protocol used to produce the MT data is known to introduce approximately 1-2 % of *E. coli* genomic DNA into the cDNA as contamination which is then reflected in the MT data. According to our analyses, 0.12 % of MG reads and 1.95 % of MT reads from this sample could be mapped onto the *E. coli* contigs which is consistent with the numbers quoted by Franzosa *et al.* (2014) [29]. This fraction of reads is sufficient for *de novo* reconstruction of approximately 70% of the *E. coli* genome. The integrative analyses of MG and MT data within IMP enables users to conveniently highlight notable cases such as this, and to further investigate inconsistencies and/or interesting characteristics within the multi-omic data.

**Fig. 5.**
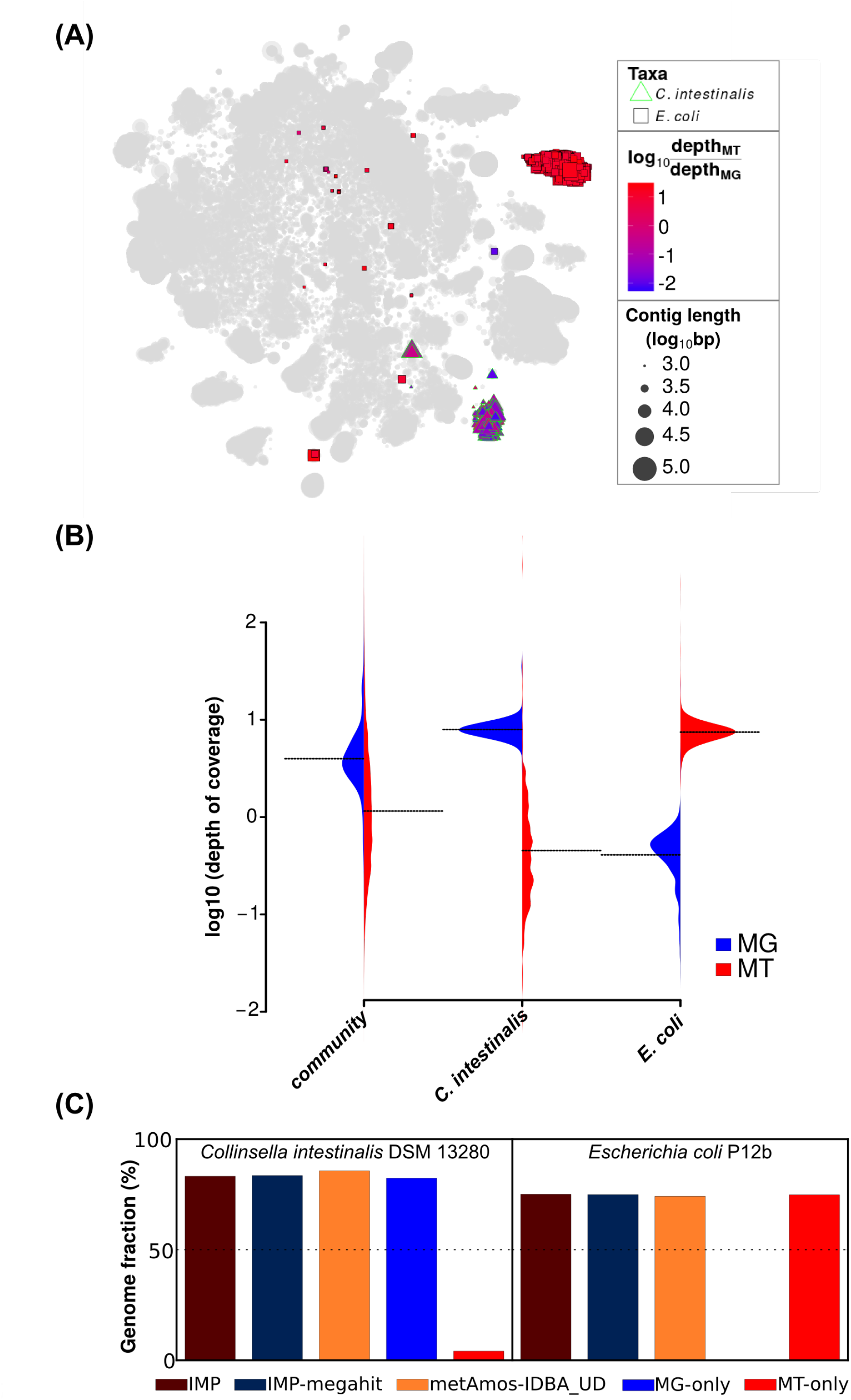
Metagenomic and metatranscriptomic data integration. **(A)** Augmented VizBin map highlighting contig subsets with sequences that are most similar to *Escherichia coli* P12b and *Colinsella intestinalis* DSM 13280 genomes. **(B)** Beanplot of metagenomic (MG) and metatranscriptomic (MT) average contig-level depth of coverage for the entire microbial community and two subsets (population-level genomes) of interest. **(C)** Recovered portion of subsets associated to the aforementioned taxa compared to a MG-only and MT-only assemblies.

## Summary

IMP was developed in order to leverage the advantages associated with integrating MG and MT data for studying microbial community structure and function *in situ* [4, 6]. Accordingly, we present a self-contained workflow which performs reproducible integrative analyses of coupled MG and MT data derived from single and unique microbial community samples. IMP encapsulates all processes including preprocessing, assembly, and analyses within an automated reproducible pipeline.

We implemented customized preprocessing and filtering procedures for MG and MT data due to the distinct nature of these different omic data types. We also evaluated the IMP-based iterative co-assembly procedure and found it to produce higher amount of output volume (a higher number of contigs and genes), thereby resulting in enhanced data usage (reflected in a higher fraction of read which can be mapped back to the contigs). IMP provides the option for the use of two state-of-the-art assemblers, whereby the default assembler, IDBA-UD, produces highly contiguous assemblies, while MEGAHIT balances the number of contigs with favorable contiguity with a high number of predicted genes and a relatively low rate of misassemblies. High quality assemblies yield better quality taxonomic information and gene annotations. Consequently, the post-assembly analysis of MG and MT data enables users to evaluate their community of interest based on taxonomy, functional potential and functional expression. The integrated co-assembly also provides the opportunity for analyses not possible based on MG data alone, such as the detection of RNA viruses and the identification of transcriptionally active populations. The output of IMP is compatible with, and may be exported to interactive tools such as VizBin [43] and Anvi’o [17] for binning and further analyses. Furthermore, the output data (annotated gene sets) may be used for the analysis and integration of additional omic data, most notably MP data.

The use of the Docker promotes reproducibility and sharing such that researchers are able to tag specific versions of the pipeline used for a particular study, thus enabling their peers to precisely replicate bioinformatic analyses workflows with relative ease and with minimal impact on overall performance of the employed bioinformatic tools [36, 37]. Static websites will be created and associated with every new version of IMP (Docker image), such that users will be able to download and launch specific versions of the pipeline to reproduce the work of others. Finally, the open source nature of IMP encourages a community-based effort to contribute and further improve the pipeline. The common scripting languages Bash, Make and Python, which Snakemake is based on [38, 39], are widely used within the bioinformatic community, thus reducing the learning curve for further development, improvement and customization by other users. Finally, the combination of open development and reproducibility should promote the general paradigm of reproducible research within the microbiome research community.

## Details and parameters of IMP

A Python (ver 3) wrapper script was implemented for user-friendly execution of IMP via the command line. The full list of dependencies, parameters (see below) and documentation are available on the IMP website (http://r3lab.uni.lu/web/imp/).

### Reproducibility

IMP is based around a Docker container that runs the Ubuntu 14.04 operating system, with all relevant dependencies. Five mounting points are defined for the Docker container with the - v option: i) input directory, ii) output directory, iii) database directory, iv) code directory, and v) configuration file directory. Environment variables are defined using the -e parameter, including: i) paired MG data, ii) paired MT data, and iii) configuration file. The latest IMP Docker image will be downloaded automatically upon launching the command, but users may also launch specific versions based on tags or use modified/customized versions of their local code base.

### Automation

Automation of the workflow is achieved using Snakemake 3.4.2, a Python-based make language implemented specifically for building reproducible bioinformatic workflows and pipelines. It allows seamless integration of Python code and shell (bash) commands, using make scripting style. It also provides checkpoints to continue interrupted analyses and/or rerun steps if required [38, 39].

### Trimming and quality filtering

Trimmomatic 0.32 [40] is used to perform trimming and quality filtering of MG and MT Illumina paired-end reads, using the following parameters: ILLUMINACLIP:TruSeq3-PE.fa:2:30:10; LEADING:20; TRAILING:20; SLIDINGWINDOW:1:3; MAXINFO:40:0.5; MINLEN:40. The parameters may be tuned in the IMP config file. The output from this step includes retained paired-ends and single-ends (mate discarded) which are all used for downstream processes.

### Ribosomal RNA filtering

SortMeRNA 2.0 is used for filtering rRNA from the MT data. The process is applied on FASTQ files for both paired- and single-end reads generated from the previous preprocessing step. Paired-end FASTQ files are interleaved prior to running SortMeRNA. If one of the mates within the paired-end read are classified as an rRNA sequence, then the entire pair is filtered out. After running SortMeRNA, the interleaved paired-end output is split into two separate paired-end FASTQ files. The filtered sequences (without rRNA) are used for the downstream processes. All available databases provided within SortMeRNA are used for filtering and the maximum memory usage parameter is set to 4 Gb (option: -m 4000).

### Read mapping

The read mapping procedure is performed using bwa mem aligner [47] with settings: -v 1 (verbose output level), -M (Picard compatibility) introducing an automated samtools header using the -R option [47]. Paired- and single-end reads are mapped separately, and the resulting alignments are merged (using samtools merge). Read mapping is performed at various steps in the workflow including: i) filtering human sequences (optional), ii) recruitment of unmapped reads within the IMP-based iterative co-assembly, and iii) mapping of preprocessed MG and MT reads to the final contig set.

### Extracting unmapped reads

The extraction of unmapped reads (paired- and single-end) begins by mapping reads to a given reference sequence (see section Read mapping). The resulting alignment file (BAM format) is used as input for the extraction of unmapped reads. A set of paired-end reads considered unmappable if both or either one of the mates do not map to the given reference. The unmapped reads are converted from BAM to FASTQ format using samtools and BEDtools 2.17.0 - bamToFastq utility [48]. Similarly, unmapped single-end reads are also extracted from the alignment information.

### Filtering host sequences

The human sequence filtering is performed by mapping the both paired- and single-end reads (see section Read mapping) onto the human genome version 38 (http://www.ncbi.nlm.nih.gov/projects/genome/assembly/grc/), followed by extraction of unmapped reads (see section Extracting unmapped reads for details). This filtering step may be omitted from the workflow using the IMP user configuration file, while users may replace the human genome with other FASTA sequences from other hosts based on their screening requirements.

### Parameters of the IMP-based iterative co-assembly

The IMP-based iterative co-assembly implements MEGAHIT 1.0.3 [16] as the MT assembler while IDBA-UD 1.1.1 [15] is used as the default co-assembler (MG + MT), with MEGAHIT [16] as an alternative option for the co-assembler. All *de novo* assemblies are performed on *k*mers ranging from 25-mers to 99-mers, with an incremental step of four. Accordingly, the command line parameters for IDBA-UD are --mink 25 --maxk 99 --step 4 --similar 0.98 --pre-correction [15]. Similarly, the command line options for MEGAHIT are --k-min 25 --k-max 99 --k-step 4, except for the MT assemblies which are performed with an additional --no-bubble option to prevent merging of bubbles within the assembly graph [16]. Furthermore, contigs generated from the MT assembly (“pseudo long-reads”) are used as input within the -l flag of IDBA-UD or -r flag of MEGAHIT [15, 16]. The parameters for the assemblies may be adjusted in the user configuration file. *Kmer* ranges may be customized in the IMP user configuration file.

### Annotation and assembly quality assessment

Prokka 1.11 [42] with the --metagenome setting is used to perform functional annotation. The default BLAST and HMM databased of Prokka are used for the functional annotation.

MetaQUAST 3.1 [41] is used to perform taxonomic annotation of contigs with the maximum number of reference genomes set to 75 (--max-ref-number 75). In addition, MetaQUAST provides various assembly statistics.

### Depth of coverage

Contig- and gene-wise depth of coverage values are calculated (per base) using BEDtools 2.17.0 [48] and aggregated (by average) using awk, adapted from the CONCOCT code [27] (script: map-bowtie2-markduplicates.sh, Github URL: https://github.com/BinPro/CONCOCT).

### Variant calling

The variant calling procedure is performed using the following tools: i) Samtools mpileup 0.1.19 [49], ii) Freebayes 0.9.21 [50] and iii) Platypus 0.8.1 [51], each using their respective default settings. The input is the merged paired- and single-end read alignment (BAM) against the final assembly FASTA file (see section Read mapping). The output files from the three methods are indexed using tabix and compressed using gzip. No filtering is applied to the variant calls, so that users may access all the information and filter them according to their requirements. The output from samtools mpileup is used for the VizBin-based visualizations.

### Non-linear dimensionality reduction of genomic signatures (NLDR-GS)

VizBin [43] performs NLDR-GS onto contigs ≥ 1kb, using default settings, to obtain 2D embeddings. Parameters can be modified in the IMP config file.

### Visualization and reporting

IMP compiles the multiple summaries and visualizations into a HTML report. FASTQC [52] is used to visualize the quality and quantity of reads before and after preprocessing. MetaQUAST [41] is used to report assembly quality and taxonomic associations of contigs. A custom script is used to generate KEGG-based [53] functional Krona plots by running KronaTools [54] (script: genes.to.kronaTable.py, GitHub URL: https://github.com/EnvGen/metagenomics-workshop). Additionally, VizBin output is integrated with the information derived from the IMP analyses, using a custom R script for analysis and visualization of the augmented maps. The R workspace (Rdat) is saved such that users are able to access it for further analyses. All the steps executed within an IMP run including parameters and runtimes are summarized in the form of a workflow diagram and a log-file.

### Output

The output generated by IMP includes a multitude of large files. Paired- and single-end FASTQ files of preprocessed MG and MT reads are provided such that the user may employ them for additional downstream analyses. The output of the IMP-based iterative co-assembly consists of a FASTA file, while the alignments/mapping of MG and MT preprocessed reads to the final co-assembly are also provided as a binary alignment format (BAM), such that users may use these for further processing. Predicted genes and their respective annotations are provided in the various formats produced by Prokka [42]. Assembly quality statistics and taxonomic annotations of contigs are provided as per the output of MetaQUAST [41]. Two-dimensional embeddings from the NLDR-GS are provided such that they can be exported to and further curated using VizBin [43] for human-augmented binning. Additionally, abundance and expression information is represented by contig- and gene-level average depth of coverage values. MG and MT genomic variant information (VCF format), including both SNPs and INDELs (insertions and deletions), is also provided.

The HTML report (see Additional file 1: HTML S1 and S2) compiles various summaries and visualizations including, i) augmented VizBin-based maps, ii) MG- and MT-level functional Krona charts [54], iii) run time information iv) detailed schematic of the steps carried out within the IMP run, v) list of parameters and commands, and vi) additional reports [FASTQC report [52], MetaQUAST report [41]]. Please refer to documentation of IMP for a detailed list and description of the output.

### Customization and further development

Basic parameters (input, output, assembler, configuration file) may be changed via the IMP command line, while more advanced parameters may be changed by editing the user configuration file (JSON format). Finally, users may directly edit the code of IMP to implement extensive changes to the pipeline, if required. The “--current” flag, within the IMP command line can be used to execute customized (local) versions of the code base. Finally, the IMP launcher script provides the option (flag: --enter) to launch the Docker container interactively, for development and testing purposes (described on the IMP website and documentation).

## Data and analyses

### Coupled metagenomic and metatranscriptomic datasets

The simulated MT data was obtained from the original authors [14], upon request. A complementary metagenome was simulated using the same set of 73 bacterial genomes used for the aforementioned simulated MT [14]. Simulated reads were obtained using the NeSSM MG simulator (default settings) [55].

Real data analyzed for this article included coupled MG and MT data, i.e. both datasets were obtained from the same unique sample. The published human fecal data derived from the healthy individual “X310763260” [29] was obtained from the NCBI Sequence Read Archive (metagenome SRA: SRX247379, metatranscriptome SRA:SRX247335). The wastewater sludge data was obtained in-house, but is available on the NCBI SRA (metagenome SRA: SRX389533, metatranscriptome SRA: SRX389534) [32].

### Iterative single-omic assemblies

In order to determine the opportune number of iterations within the IMP-based iterative co-assembly strategy within IMP, we first performed an initial assembly using IMP preprocessed SM, HF and WW MG reads with IDBA-UD [15] together with cap3 [56], which was used to further collapse the contigs and reduce the redundancy of the assembly. This initial assembly was followed by a total of three assembly iterations. Each iteration was made up of four separate steps: i) extraction of reads unmappable to the previous assembly (using the procedure explained in Extracting unmapped reads), ii) assembly of unmapped reads using IDBA-UD [15], iii) merging/collapsing the contigs from the previous assembly using cap3 [56], and iv) evaluation of the merged assembly using MetaQUAST [41]. The assembly was evaluated in terms of the per-iteration increase in mappable reads, assembly length, numbers of contigs ≥ 1 kb, and number of unique genes.

Similar iterative assemblies were also performed for MT data of SM, HF and WW using MEGAHIT [16] except, CD-HIT-EST [57] was used to collapse the contigs at ≥ 95% identity (-c 0.95) while MetaGeneMark [58] was used to predict genes. The parameters and settings of the other programs were the same as those defined in Details and parameters of IMP.

The contigs from the first iteration of both the MG and MT iterative assemblies were selected to represent the control single-omic (MG-only and MT-only) assemblies and were compared against co-assemblies.

### Execution of pipelines

MetAMOS was executed on each dataset using: i) the default setting, with SOAPdenovo as assembler (using a length of 31-mers), and ii) a custom version with IDBA-UD as the assembler (option: -a idba-ud) using both MG and MT paired-end FASTQ reads as input. All computations using MetAMOS were set to use eight computing cores per run (option: -p 8). Similarly, IMP was executed for each dataset using different assemblers for the co-assembly step: i) default setting using IDBA-UD, and ii) MEGAHIT (option: -a megahit). Additionally, the analysis of HF data included the preprocessing step of filtering human genome sequences, which was omitted for WW data. Illumina TruSeq2 adapter trimming was used for WW data preprocessing, since the information was available. Computation was performed using eight computing cores. The customized parameters were specified in the IMP configuration file (see Additional file 1: HTML S1 and S2 for exact configurations).

### Assembly assessment and comparison

Assemblies were assessed and compared at the contig level (scaffolds not considered), using MetaQUAST [41]. The gene calling function (flag: -f) was utilized to obtain the number of genes which were predicted from the various assemblies. An additional parameter within MetaQUAST was used for ground truth assessment of the simulated mock (SM) community assemblies, by providing the list of 73 reference genomes (flag: -R). MetaQUAST was applied to compare: i) single-omic assemblies and multi-omic co-assemblies and ii) co-assemblies from different pipelines.

### Analysis of contigs assembled from MT data

A list of contigs with no MG depth of coverage together with additional information on these (contig length, annotation, MT depth of coverage) was retrieved using the R workspace (Rdat) which is provided as part IMP output. The sequences of these contigs were extracted and subjected to a BLAST search on NCBI to determine their potential origin. Furthermore, contigs with length ≥ 1kb, average depth of coverage ≥ 20 bases and containing genes encoding known virus/bacteriophage functions were extracted.

### Analysis of subsets of contigs

Subsets of contigs were identified by visual inspection of augmented VizBin maps generated by IMP. Detailed inspection of contig-level MT to MG depth of coverage ratios was carried out using the R workspace provided as part of IMP output. The alignment information of contigs to isolate genomes provided by MetaQUAST [41] were used to highlight subsets of contigs aligning to genomes of *Escherichia coli* P12B strain (*E. coli*) and *Collinsella intestinalis* DSM 13280 **(C.** *intestinalis).*

MetaQUAST [41] was used to compare the three co-assemblies carried out on the HF dataset (using IMP, IMP -MEGAHIT and MetAMOS-IDBA-UD) against the corresponding single-omic MG and MT assemblies (see section Iterative single-omic assemblies). For the HF data, corresponding reference genomes were extracted from the IMP output and were provided to MetaQUAST (flag: -R) as the reference genome set.

### Computational platforms

IMP and MetAMOS were executed on a Dell R820 machine with 32 Intel(R) Xeon(R) CPU E5-4640 @ 2.40GHz physical computing cores (64 virtual), 1024 TB of DDR3 RAM (32 GB per core) with Debian 7 Wheezy as the operating system. Additional computations outside the scope of the pipelines (IMP and MetAMOS) were performed on the Gaia cluster of the University of Luxembourg HPC platform [59].

## Availability

IMP software and code are available under the BSD-4-Clause license, on the LCSB R3 website: http://r3lab.uni.lu/web/imp/. Scripts and commands for additional analyses are available at: https://git-r3lab.uni.lu/shaman.narayanasamy/IMP_article_analyses.

## Competing interests

The authors declare that they have no competing interests

## Authors’ contributions

SN, NP, EELM, PM, PW conceived the analysis and designed the workflow. SN, YJ, MH, CCL developed the software, wrote the documentation and tested the software. YJ ensured reproducibility of the software. SN performed data analyses. EELM, PM, AHB, AK, NP, PW participated in discussions and tested the software. SN, EELM, AHB, PM, NP, AK, MH, PW wrote and edited the manuscript. PW designed and supported the project. All authors read and agreed on the final version of the manuscript.

## Abbreviations

NGS: next-generation sequencing
Contigs: contiguous sequence(s)
cDNA: complementary-DNA
MG: Metagenomic
MT: Metatranscriptomic
MP: Metaproteome/metaproteomic
SNPs: single nucleotide polymorphisms
INDELs: insertions and deletions
rRNA: ribosomal RNA
IMP: Integrated Meta-omic Pipeline
SM: Simulated mock
HF: Human fecal
WW: Wastewater
bp: base pair
Kb: kilo base
KEGG: Kyoto Encyclopedia of Genes and Genomes
VCF: Variant call format
SRA: Sequence read archive
NCBI: National Center for Biotechnology Information

## Acknowledgements

We would like to acknowledge John Larsson from SciLifeLab (Sweden) for kindly providing the KEGG-based functional Krona plot scripts. Albi Celaj from the University of Toronto is thanked for supplying the *in silico* simulated metatranscriptomic data and the corresponding reference genomes. The University of Luxembourg High Performance Computing (HPC) facility is duly thanked for providing and maintaining the computing platform. The Reproducible Research Results (R3) team of the Luxembourg Centre for Systems Biomedicine is acknowledged for support of the project and for promoting reproducible research. This work was supported by an ATTRACT program grant (A09/03), a European Union Joint Programming in Neurodegenerative Diseases grant (INTER/JPND/12/01) and a proof of concept grant (PoC/13/02) to PW, Aide à la Formation Recherche (AFR) grants to SN (PHD-2014-1/7934898), and a CORE junior (C15/SR/10404839) to EELM, all funded by the Luxembourg National Research Fund (FNR).

## Additional files

The following additional data are available with the online version of this article.

### Additional file 1

File format: HTML

Title of data: Supplementary IMP HTML reports.

Description of data: HTML S1 and S2 are reports produced by IMP for the analysis of the human fecal (HF) microbial community and wastewater (WW) sludge microbial community datasets.

### Additional file 2

File format: PDF

Title of data: Supplementary figures.

Description of data: Figures S1 to S4. Detailed figure legends available within file.

### Additional file 3

File format: MS Excel (XLSX)

Title of data: Supplementary tables.

Description of data: Tables S1 to S9. Detailed table legends available within file.

## References

1. Turnbaugh PJ, Ley RE, Hamady M, Fraser-liggett C, Knight R, Gordon JI: The human microbiome project: exploring the microbial part of ourselves in a changing world. Nature 2007, 449:804–810.

2. Rittmann BE: Microbial ecology to manage processes in environmental biotechnology. Trends Biotechnol 2006, 24:261–266.

3. Stewart EJ: Growing unculturable bacteria. J Bacteriol 2012, 194:4151–4160.

4. Narayanasamy S, Muller EEL, Sheik AR, Wilmes P: Integrated omics for the identification of key functionalities in biological wastewater treatment microbial communities. Microb Biotechnol 2015, 8:363–368.

5. Segata N, Waldron L, Ballarini A, Narasimhan V, Jousson O, Huttenhower C, Boernigen D, Tickle TL, Morgan XC, Garrett WS, Huttenhower C: Computational meta’omics for microbial community studies. Mol Syst Biol 2013, 9:666.

6. Muller EEL, Glaab E, May P, Vlassis N, Wilmes P: Condensing the omics fog of microbial communities. Trends Microbiol 2013, 21:325–333.

7. Roume H, Muller EEL, Cordes T, Renaut J, Hiller K, Wilmes P: A biomolecular isolation framework for eco-systems biology. ISME J 2013, 7:110–121.

8. Roume H, Heintz-Buschart A, Muller EEL, Wilmes P: Sequential isolation of metabolites, RNA, DNA, and proteins from the same unique sample. Methods Enzymol 2013, 531:219–236.

9. Solomon K V, Haitjema CH, Thompson DA, O’Malley MA: Extracting data from the muck: deriving biological insight from complex microbial communities and non-model organisms with next generation sequencing. Curr Opin Biotechnol 2014, 28C:103–110.

10. Sunagawa S, Mende DR, Zeller G, Izquierdo-Carrasco F, Berger SA, Kultima JR, Coelho LP, Arumugam M, Tap J, Nielsen HB, Rasmussen S, Brunak S, Pedersen O, Guarner F, de Vos WM, Wang J, Li J, Doré J, Ehrlich SD, Stamatakis A, Bork P: Metagenomic species profiling using universal phylogenetic marker genes. Nat Methods 2013, 10:1196–1199.

11. Li J, Jia H, Cai X, Zhong H, Feng Q, Sunagawa S, Arumugam M, Kultima JR, Prifti E, Nielsen T, Juncker AS, Manichanh C, Chen B, Zhang W, Levenez F, Wang J, Xu X, Xiao L, Liang S, Zhang D, Zhang Z, Chen W, Zhao H, Al-Aama JY, Edris S, Yang H, Wang J, Hansen T, Nielsen HB, Brunak S, et al.: An integrated catalog of reference genes in the human gut microbiome. Nat Biotechnol 2014, 32:834–841.

12. Treangen TJ, Koren S, Sommer DD, Liu B, Astrovskaya I, Ondov B, Darling AE, Phillippy AM, Pop M: MetAMOS: a modular and open source metagenomic assembly and analysis pipeline. Genome Biol 2013, 14:R2.

13. Nalbantoglu OU, Way SF, Hinrichs SH, Sayood K: RAIphy: phylogenetic classification of metagenomics samples using iterative refinement of relative abundance index profiles. BMC Bioinformatics 2011, 12:41.

14. Celaj A, Markle J, Danska J, Parkinson J: Comparison of assembly algorithms for improving rate of metatranscriptomic functional annotation. Microbiome 2014, 2:39.

15. Peng Y, Leung HCM, Yiu SM, Chin FYL: IDBA-UD: a de novo assembler for single-cell and metagenomic sequencing data with highly uneven depth. Bioinformatics 2012, 28:1420–1428.

16. Li D, Liu C-M, Luo R, Sadakane K, Lam T-W: MEGAHIT: an ultra-fast single-node solution for large and complex metagenomics assembly via succinct de Bruijn graph. Bioinformatics 2015, 31:1674–1676.

17. Eren AM, Esen ÖC, Quince C, Vineis JH, Morrison HG, Sogin ML, Delmont TO: Anvi’o: an advanced analysis and visualization platform for ‘omics data. PeerJ 2015, 3:e1319.

18. Leimena MM, Ramiro-Garcia J, Davids M, van den Bogert B, Smidt H, Smid EJ, Boekhorst J, Zoetendal EG, Schaap PJ, Kleerebezem M: A comprehensive metatranscriptome analysis pipeline and its validation using human small intestine microbiota datasets. BMC Genomics 2013, 14:530.

19. Leung HCM, Yiu S-M, Parkinson J, Chin FYL: IDBA-MT: de novo assembler for metatranscriptomic data generated from next-generation sequencing technology. J Comput Biol 2013, 20:540–550.

20. Grabherr MG, Haas BJ, Yassour M, Levin JZ, Thompson DA, Amit I, Adiconis X, Fan L, Raychowdhury R, Zeng Q, Chen Z, Mauceli E, Hacohen N, Gnirke A, Rhind N, Palma F, Birren BW, Nusbaum C, Lindblad-toh K, Friedman N, Regev A: Full-length transcriptome assembly from RNA-Seq data without a reference genome. Nat Biotechnol 2011, 29:644–652.

21. Kultima JR, Sunagawa S, Li J, Chen W, Chen H, Mende DR, Arumugam M, Pan Q, Liu B, Qin J, Wang J, Bork P: MOCAT: a metagenomics assembly and gene prediction toolkit. PLoS One 2012, 7:e47656.

22. Luo R, Liu B, Xie Y, Li Z, Huang W, Yuan J, He G, Chen Y, Pan Q, Liu Y, Tang J, Wu G, Zhang H, Shi Y, Liu Y, Yu C, Wang B, Lu Y, Han C, Cheung DW, Yiu S-M, Peng S, Xiaoqian Z, Liu G, Liao X, Li Y, Yang H, Wang J, Lam T-W, Wang J: SOAPdenovo2: an empirically improved memory-efficient short-read de novo assembler. Gigascience 2012, 1:18.

23. Lai B, Wang F, Wang X, Duan L, Zhu H: InteMAP: Integrated metagenomic assembly pipeline for NGS short reads. BMC Bioinformatics 2015, 16:244.

24. Laczny CC, Pinel N, Vlassis N, Wilmes P: Alignment-free visualization of metagenomic data by nonlinear dimension reduction. Sci Rep 2014, 4:4516.

25. Albertsen M, Hugenholtz P, Skarshewski A, Nielsen KL, Tyson GW, Nielsen PH: Genome sequences of rare, uncultured bacteria obtained by differential coverage binning of multiple metagenomes. Nat Biotechnol 2013, 31:533–538.

26. Nielsen HB, Almeida M, Juncker AS, Rasmussen S, Li J, Sunagawa S, Plichta DR, Gautier L, Pedersen AG, Le Chatelier E, Pelletier E, Bonde I, Nielsen T, Manichanh C, Arumugam M, Batto J-M, Quintanilha Dos Santos MB, Blom N, Borruel N, Burgdorf KS, Boumezbeur F, Casellas F, Doré J, Dworzynski P, Guarner F, Hansen T, Hildebrand F, Kaas RS, Kennedy S, Kristiansen K, et al.: Identification and assembly of genomes and genetic elements in complex metagenomic samples without using reference genomes. Nat Biotechnol 2014, 32:822–828.

27. Alneberg J, Bjarnason BS, de Bruijn I, Schirmer M, Quick J, Ijaz UZ, Lahti L, Loman NJ, Andersson AF, Quince C: Binning metagenomic contigs by coverage and composition. Nat Methods 2014, 11:1144–1146.

28. Kang DD, Froula J, Egan R, Wang Z: MetaBAT, an efficient tool for accurately reconstructing single genomes from complex microbial communities. Peer J 2015, 3:e1165.

29. Franzosa E a, Morgan XC, Segata N, Waldron L, Reyes J, Earl AM, Giannoukos G, Boylan MR, Ciulla D, Gevers D, Izard J, Garrett WS, Chan AT, Huttenhower C: Relating the metatranscriptome and metagenome of the human gut. Proc Natl Acad Sci U S A 2014, 111:E2329–E2338.

30. Hultman J, Waldrop MP, Mackelprang R, David MM, Mcfarland J, Blazewicz SJ, Harden J, Turetsky MR, Mcguire AD, Shah MB, Verberkmoes NC, Lee LH: Multi-omics of permafrost, active layer and thermokarst bog soil microbiomes. Nature 2015, 521:208–2112.

31. Bremges A, Maus I, Belmann P, Eikmeyer F, Winkler A, Albersmeier A, Pühler A, Schlüter A, Sczyrba A: Deeply sequenced metagenome and metatranscriptome of a biogas-producing microbial community from an agricultural production-scale biogas plant. Gigascience 2015, 4:33.

32. Muller EEL, Pinel N, Laczny CC, Hoopman MR, Narayanasamy S, Lebrun LA, Roume H, Lin J, May P, Hicks ND, Heintz-Buschart A, Wampach L, Liu CM, Price LB, Gillece JD, Guignard C, Schupp JM, Vlassis N, Baliga NS, Moritz RL, Keim PS, Wilmes P: Community integrated omics links the dominance of a microbial generalist to fine-tuned resource usage. Nat Commun 2014, 5:5603.

33. Roume H, Heintz-Buschart A, Muller EEL, May P, Satagopam VP, Laczny CC, Narayanasamy S, Lebrun LA, Hoopmann MR, Schupp JM, Gillece JD, Hicks ND, Engelthaler DM, Sauter T, Keim PS, Moritz RL, Wilmes P: Comparative integrated omics: identification of key functionalities in microbial community-wide metabolic networks. npj Biofilms Microbiomes 2015, 1:15007.

34. Vanwonterghem I, Jensen PD, Ho DP, Batstone DJ, Tyson GW: Linking microbial community structure, interactions and function in anaerobic digesters using new molecular techniques. Curr Opin Biotechnol 2014, 27:55–64.

35. Kenall A, Edmunds S, Goodman L, Bal L, Flintoft L, Shanahan DR, Shipley T: Better reporting for better research: a checklist for reproducibility. BMC Neurosci 2015, 16:44.

36. Belmann P, Dröge J, Bremges A, McHardy AC, Sczyrba A, Barton MD: Bioboxes: standardised containers for interchangeable bioinformatics software. Gigascience 2015, 4:47.

37. Di Tommaso P, Palumbo E, Chatzou M, Prieto P, Heuer ML, Notredame C: The impact of Docker containers on the performance of genomic pipelines. PeerJ 2015, 3:e1273.

38. Köster J, Rahmann S: Snakemake-a scalable bioinformatics workflow engine. Bioinformatics 2012, 28:2520–2522.

39. Koster J: Reproducibility in next-generation sequencing analysis. http://dx.doi.org/10.17877/DE290R-7242 (2014). Accessed 05 Feb 2015.

40. Bolger AM, Lohse M, Usadel B: Trimmomatic: a flexible trimmer for Illumina sequence data. Bioinformatics 2014, 30:2114–2120.

41. Mikheenko A, Saveliev V, Gurevich A: MetaQUAST: evaluation of metagenome assemblies. Bioinformatics 2015:btv697.

42. Seemann T: Prokka: rapid prokaryotic genome annotation. Bioinformatics 2014, 30:2068–2069.

43. Laczny CC, Sternal T, Plugaru V, Gawron P, Atashpendar A, Margossian HH, Coronado S, der Maaten L van, Vlassis N, Wilmes P: VizBin - an application for reference-independent visualization and human-augmented binning of metagenomic data. Microbiome 2015, 3:1.

44. Mende DR, Waller AS, Sunagawa S, Järvelin AI, Chan MM, Arumugam M, Raes J, Bork P: Assessment of metagenomic assembly using simulated next generation sequencing data. PLoS One 2012, 7:e31386.

45. Schürch AC, Schipper D, Bijl MA, Dau J, Beckmen KB, Schapendonk CME, Raj VS, Osterhaus ADME, Haagmans BL, Tryland M, Smits SL: Metagenomic survey for viruses in Western Arctic caribou, Alaska, through iterative assembly of taxonomic units. PLoS One 2014, 9:e105227.

46. Reyes A, Blanton L V., Cao S, Zhao G, Manary M, Trehan I, Smith MI, Wang D, Virgin HW, Rohwer F, Gordon JI: Gut DNA viromes of Malawian twins discordant for severe acute malnutrition. Proc Natl Acad Sci U S A 2015, 112:11941–11946.

47. Li H, Durbin R: Fast and accurate short read alignment with Burrows-Wheeler transform. Bioinformatics 2009, 25:589–595.

48. Quinlan AR, Hall IM: BEDTools: a flexible suite of utilities for comparing genomic features. Bioinformatics 2010, 26:841–842.

49. Li H, Handsaker B, Wysoker A, Fennell T, Ruan J, Homer N, Marth G, Abecasis G, Durbin R: The Sequence Alignment/Map format and SAMtools. Bioinformatics 2009, 25:2078–2079.

50. Garrison E, Marth G: Haplotype-based variant detection from short-read sequencing. bioRxiv 2012:9.

51. Rimmer A, Phan H, Mathieson I, Iqbal Z, Twigg SRF, Wilkie AOM, McVean G, Lunter G: Integrating mapping-, assembly- and haplotype-based approaches for calling variants in clinical sequencing applications. Nat Genet 2014, 46:912–918.

52. Patel RK, Jain M: NGS QC Toolkit: a toolkit for quality control of next generation sequencing data. PLoS One 2012, 7:e30619.

53. Kanehisa M: KEGG: Kyoto Encyclopedia of Genes and Genomes. Nucleic Acids Res 2000, 28:27–30.

54. Ondov BD, Bergman NH, Phillippy AM: Interactive metagenomic visualization in a Web browser. BMC Bioinformatics 2011, 12:385.

55. Jia B, Xuan L, Cai K, Hu Z, Ma L, Wei C: NeSSM: a Next-generation Sequencing Simulator for Metagenomics. PLoS One 2013, 8:e75448.

56. Huang X, Madan A: CAP3: A DNA sequence assembly program. Genome Res 1999, 9:868–877.

57. Fu L, Niu B, Zhu Z, Wu S, Li W: CD-HIT: accelerated for clustering the next-generation sequencing data. Bioinformatics 2012, 28:3150–3152.

58. Zhu W, Lomsadze A, Borodovsky M: Ab initio gene identification in metagenomic sequences. Nucleic Acids Res 2010, 38:e132.

59. Varrette S, Bouvry P, Cartiaux H, Georgatos F: Management of an Academic HPC Cluster : The UL Experience. Proc 2014 Int Conf High Perform Comput Simul 2014:959–967.

